# Identification of a suitable endogenous control miRNA in bone aging and senescence

**DOI:** 10.1101/2022.02.03.479003

**Authors:** Japneet Kaur, Dominik Saul, Madison L. Doolittle, Jennifer L. Rowsey, Stephanie J. Vos, Joshua N. Farr, Sundeep Khosla, David G. Monroe

**Author notes:** To whom correspondence should be addressed: David G. Monroe, PhD, Department of Medicine, Division of Endocrinology, Mayo Clinic College of Medicine, Guggenheim 7-11A, 200 First Street SW, Rochester, MN 55905, USA. Disclosure: The authors have nothing to disclose.

## Abstract

**Objective:** MicroRNAs (miRNAs) are promising tools as biomarkers and therapeutic agents in various chronic diseases such as osteoporosis, cancers, type I and II diabetes, and cardiovascular diseases. Considering the rising interest in the regulatory role of miRNAs in bone metabolism, aging, and cellular senescence, accurate normalization of qPCR-based miRNA expression data using an optimal endogenous control becomes crucial.

**Methods:** We used a systematic approach to select candidate endogenous control miRNAs that exhibit high stability with aging from our miRNA sequence data and literature search. Validation of miRNA expression was performed using qPCR and their comprehensive stability was assessed using the RefFinder tool which is based on four statistical algorithms: GeNorm, NormFinder, BestKeeper, and comparative delta CT. The selected endogenous control was then validated for its stability in mice and human bone tissues, and in bone marrow stromal cells (BMSCs) following induction of senescence and senolytic treatment. Finally, the utility of selected endogenous control versus *U6* was tested by using each as a normalizer to measure the expression of *miR-34a*, a miRNA known to increase with age and senescence.

**Results:** Our results show that *Let-7f* did not change across the groups with aging, senescence or senolytic treatment, and was the most stable miRNA, whereas *U6* was the least stable. Moreover, using *Let-7f* as a normalizer resulted in significantly increased expression of *miR-34a* with aging and senescence and decreased expression following senolytic treatment. However, the expression pattern for *miR-34a* reversed for each of these conditions when *U6* was used as a normalizer.

**Conclusions:** We show that optimal endogenous control miRNAs, such as *Let-7f*, are essential for accurate normalization of miRNA expression data to increase the reliability of results and prevent misinterpretation. Moreover, we present a systematic strategy that is transferrable and can easily be used to identify endogenous control miRNAs in other biological systems and conditions.

## INTRODUCTION

MicroRNAs (miRNAs) are a class of evolutionarily conserved short non-coding RNAs (∼22 nucleotides (nt)) that bind to the 3’-untranslated regions (3’-UTR) of target mRNAs, resulting in mRNA degradation and suppression of translation (Bartel, 2004; Bushati & Cohen, 2007). Following the discovery of *lin-4*, the first miRNA to be discovered that repressed *lin-14* message in *C. elegans*, thousands of miRNAs have been identified in various plants and animal species including humans that regulate a wide range of biological functions such as cell proliferation, apoptosis, regeneration, and metabolism (Bartel, 2004; Lee et al., 1993; Wightman et al., 1993). Moreover, their ability to circulate throughout the body in cell-free encapsulated vesicles called extracellular vesicles or exosomes (30-150nm-diameter) have made them potent signaling molecules, disease biomarkers, and essential prognostic and therapeutic tools in chronic age-related diseases such as osteoporosis, cardiovascular diseases, type I and II diabetes, and cancers (Bottani et al., 2019; Condrat et al., 2020; Foessl et al., 2019; Januszewski et al., 2021; Mandourah et al., 2018; Margaritis et al., 2021; Materozzi et al., 2018).

Aging is a major risk factor for most chronic diseases and is characterized by progressive multi-organ deterioration and tissue dysfunction, and long-term accumulation of permanently growth-arrested ‘senescent cells’ in various tissues (Franceschi et al., 2018; Jaul & Barron, 2017; Kaur et al., 2019; Kaur & Farr, 2020; Khosla et al., 2020; McHugh & Gil, 2018). Originally described by Hayflick in 1961, cellular senescence is a cell fate that results in irreversible cell cycle arrest in response to a stressor, such as DNA damage, oncogenic insult, and reactive oxygen species, etc., along with activation of tumor suppressor pathways (mainly the ATM/p53/ p21^CIP1^ and p16^INK4a^/retinoblastoma pathways), changes in chromatin organization, resistance to apoptosis, and excessive production of pro-inflammatory cytokines (the senescence-associated secretory phenotype (SASP)) that induce senescence in neighboring healthy cells. Senescent cells are characterized by their flattened morphology and increased senescence-associated β-galactosidase (SA-β-Gal) activity and mRNA expression of *p16*^*ink4a*^ and *p21*^*Cip1*^ tumor suppressor genes (Baker et al., 2011; Khosla et al., 2020; Kirkland & Tchkonia, 2017). Age-related accumulation of senescent cells is one of the main causative factors of osteoporosis that can be impeded by selectively targeting these cells using senolytic drugs such as a combination of dasatinib plus quercetin (D+Q) that specifically eliminates senescent cells via senolysis (seno = senescence; lysis = kill) to thereby extend healthspan, and result in decreased number of SA-β-Gal positive cells and reduction in *p16*^*Ink4a*^ and *p21*^*Cip1*^ gene expression (Farr et al., 2017; Zhu et al., 2015). Therefore, there is mounting interest in developing senolytics for alleviating age-related tissue dysfunction and to identify appropriate techniques to monitor senolysis in various biological settings.

Recently, miRNAs have emerged as key regulators of bone remodeling, aging, and cellular senescence, and are known to be mechanistically involved in regulating these conditions (Grillari et al., 2021; Potter et al., 2021; Thalyana & Slack, 2012). Despite the rapid increase in the availability of high-throughput deep sequencing for characterization of miRNA transcriptome, qPCR remains the gold-standard and most widely used method for measuring miRNA expression (Donati et al., 2019; Drobna et al., 2018). However, the accuracy of their expression data obtained via qPCR is highly dependent on the choice of an appropriate endogenous normalizer or control. MicroRNA endogenous controls need to be assessed for each experiment separately since their expression varies depending on their tissue of origin and the disease state or condition being tested (Landgraf et al., 2007). Generally, a suitable endogenous control is a gene or a combination of genes that is structurally similar to the genes being tested, is highly stable, and is relatively abundantly expressed across the different models and conditions being tested (Drobna et al., 2018). Most of the studies investigating miRNAs use small nuclear RNAs (snRNAs) such as *U6* as their endogenous normalizer (Beyer et al., 2015; He et al., 2013; H. Li et al., 2018; T. Xu et al., 2020).

However, the structural differences between snRNAs (150nt) and miRNAs (20-24nt) can impact their isolation yield, reverse transcription, and amplification efficiencies, leading to inaccurate and unreliable results and possible misinterpretation of the mechanistic and therapeutic roles of the miRNAs being tested (Drobna et al., 2018).

With rising interest in the role of miRNAs in bone aging and cellular senescence, a pertinent question is which miRNA will serve as an optimal endogenous normalizer. To answer this question, we used a stepwise approach to identify a suitable endogenous control miRNA from miRNA sequence data and existing literature that remains stable with aging in bone tissue of mice and humans and with senescence as well as in response to senolytic treatment in culture. Following this, the applicability of the selected miRNA control versus *U6* was tested by using each as a normalizer to measure the expression of *miR- 34a-5p*, a miRNA well known in the literature to increase with aging and senescence in various tissues and cells and predicted to target *p16*^*ink4a*^ and *p21*^*Cip1*^ in humans as per our bioinformatic analyses (Fulzele et al., 2019; Q. Li et al., 2019; Zhang et al., 2018). The expression of *miR-34a* was evaluated in bone biopsies from young and old women, previously shown to have increased expression of the senescence-related genes *p16*^*ink4a*^, *p21*^*Cip1*^, and a subset of SASP factors, and in mouse BMSCs following induction of senescence and senolytic treatment (Farr et al., 2016). This study was driven by our observation on the lack of stability of *U6* across different biological conditions and its widespread use as an endogenous normalizer in the absence of an optimal endogenous control miRNA.

## Materials and Methods

### Bone tissue samples (mouse and human)

All mouse studies were conducted in accordance with NIH guidelines and approved by the Institutional Animal Care and Use Committee at the Mayo Clinic. The vertebrae isolated from both 6-month-old and 24- month-old male C57BL/6N mice (n = 10/group) were cleaned of associated muscle and connective tissues, minced into ∼1-mm pieces and sequentially digested twice for 30-min in endotoxin-free collagenase (Liberase; Roche Diagnostics GmbH, Mannheim, Germany) to obtain osteocyte-enriched bone samples. These were then homogenized in QIAzol reagent (Qiagen, Valencia, CA) for total RNA isolation and used for miRNA sequencing. (Farr et al., 2016).

Similarly, vertebrae from young (6-month) and old (24-month) male and female mice prepared in the same way were used for RT-qPCR analyses. The metaphysis and diaphysis were also cleaned of the muscle and connective tissue and homogenized in QIAzol reagent (Qiagen). The human bone biopsy samples used in this study were as described in previous publications (Farr et al., 2015, 2016; Fujita et al., 2014); they were small needle bone biopsies isolated from the posterior iliac crest procured from 10 young premenopausal women (mean age ± SD, 27 ± 3 years; range 23 to 30 years) and 10 old postmenopausal women (78 ± 5 years; range 72 to 87 years). Postmenopausal status was established by the absence of menses for >1 year and serum follicle stimulate hormone levels >20 IU/L. Extensive exclusion criteria for these patients and the study protocol are as previously described (Farr et al., 2015). The RNA from these bone biopsy samples was reanalyzed for miRNA expression. All human studies were approved by the Mayo Clinic Institutional Review Board.

### Cell culture

Bone marrow stromal cells (BMSCs) were isolated from 3-month-old and 33-month-old mice C57BL/6N mice after removing the epiphyseal growth plates from the tibias and femurs and flushed with Dulbecco’s Modified Eagle Medium (DMEM, ThermoFisher Scientific, Waltham, MA, USA) supplemented with 1 × antibiotic/antimycotic (ThermoFisher Scientific), 1 × Glutamax, and 15% (v/v) fetal bovine serum (GE Healthcare Life Sciences HyClone Laboratories, Logan, UT). Half media change was done on day 3 and the cells were plated for the experiment on day 7 at a cell density of 10^4^ cells/cm^2^ in 12-well tissue culture plates and allowed to grow to 70-80% confluence. To induce senescence, the cells from the 3- month-old mice were then treated with 20uM of etoposide (MilliporeSigma, St. Lous, MO) or vehicle (0.1 % DMSO) for 48 hours followed by maintenance in growth media for 6 days. For senolytic treatment, cells from 33-month-old mice were treated with 0.2□µM dasatinib (LC Laboratories, Woburn, MA) and 20□µM quercetin (Sigma-Aldrich, St. Louis, MO) or vehicle (DMSO) for 24□h, washed, and allowed to recover for 1 additional day (Zhou et al., 2021). Cells were lysed in QIAzol reagent (Qiagen) for RNA isolation or fixed with 4% paraformaldehyde (PFA) for SA-β-Gal assay.

### SA-β-Gal Assay

Cellular SA-β-Gal activity was assayed as previously described (M. Xu et al., 2015). In brief, following fixation, the BMSCs were washed three times with 1x PBS before being incubated in SA- β-Gal activity solution (pH 6.0) at 37 °C for 16–18 hours. The enzymatic reaction was stopped by washing cells or tissues three to five times with ice-cold 1x PBS.

### miRNA-sequencing

miRNA sequence analysis was performed on the osteocyte-enriched bone samples obtained from collagenase digested vertebrae of young (n=10, 6-month) and old (n=10, 24-month) C57BL/6N male mice using previously described methods (Farr et al., 2016). MicroRNAs with raw read counts less than 4 in both the groups were classified as unreliable and deleted. NormFinder analyses was done using R (4.0.1) to assess the stability scores of miRNA expression across the groups and then the miRNAs were sorted to select the ones with the highest normalized counts. The predicted target mRNAs for *miR-34a-5p* in humans were determined using miRNet (https://www.mirnet.ca/miRNet/home.xhtml).

### Analyses of RT-qPCR expression stability

For assessing the expression stability of the RT-qPCR data, the median CT values (from triplicates) collected for each candidate endogenous control miRNA for each sample were averaged and used as the input data in RefFinder tool (https://www.heartcure.com.au/reffinder/) which generated a comprehensive stability ranking and individual ranking for each of the four algorithms: GeNorm (Vandesompele et al., 2002), NormFinder (Andersen et al., 2004), BestKeeper (Pfaffl et al., 2004), and comparative delta CT (Silver et al., 2006).

### qPCR analyses of mRNA and miRNA assays

Total RNA (125ng) was used to generate cDNA using the High-Capacity cDNA Reverse Transcription Kit (Applied Biosystems, Carlsbad, CA) according to the manufacturer’s instructions. qPCR analysis was performed using the ABI Prism 7900HT Real-Time System instrument (Applied Biosystems) with SYBR Green reagent (Qiagen) for mRNA and as previously described (Farr et al., 2015). Data normalization was performed based on 5 reference genes (*Actb, Gapdh, Polr2a, Rpl13a, Tuba1a*) depending on their stability and threshold calculations are as previously described (I. Mödder et al., 2011). The oligonucleotide sequences for the genes measured in this study were designed using the Primer Express program (Applied Biosystems) and are available upon request. For the miRNA expression data, total RNA (30ng) including the miRNA fraction was reverse transcribed using the miRCURY LNA RT Kit (Qiagen) as per manufacturer’s protocol. The individual miRNA assays used in this study were purchased from Qiagen and used with the miRCURY LNA miRNA PCR Starter Kit (Qiagen) according to the manufacturer’s instructions. Data normalization was performed using the *Let-7f* and *RNU6B (U6)* small nucleolar RNA (snRNA) as specified in Results.

### Data analysis

miRNA expression levels of the candidate endogenous controls were determined by CT values measured in triplicates. The fold-changes were determined using the relative quantification method (2^−ΔΔCt^) with the selected endogenous control. All generated quantitative data was presented as the mean ± SE (unless otherwise specified) and differences were analyzed using t-test analysis with *p*<0.05 (two-tailed) considered to be statistically significant. Statistical analyses were performed using the Statistical Package for the Social Sciences for Windows, Version 25.0 (SPSS, Chicago, IL).

## Results

### Identifying suitable endogenous controls from the miRNA sequence dataset

We utilized a systematic strategy to identify suitable endogenous controls from our miRNA sequence dataset obtained from the osteocyte-enriched bone samples of young (6-month) and old (24-month) mice (n=10/group) (Fig. 1). To select the optimal number of miRNAs from the total of 633 expressed in our dataset, we extended the approach followed by Drobna et al. and used their ‘iterative analysis of stability’ algorithm based on NormFinder to shortlist the best 100 miRNAs with the highest stability from our dataset (Drobna et al., 2018). A standard NormFinder analysis identified the most stable miRNAs and then the signals of these miRNAs were averaged with the remaining N-1 miRNAs individually generating a stability value. Generally, the addition of a miRNA is expected to decrease the stability score, thereby indicating higher stability.

**Figure 1.**
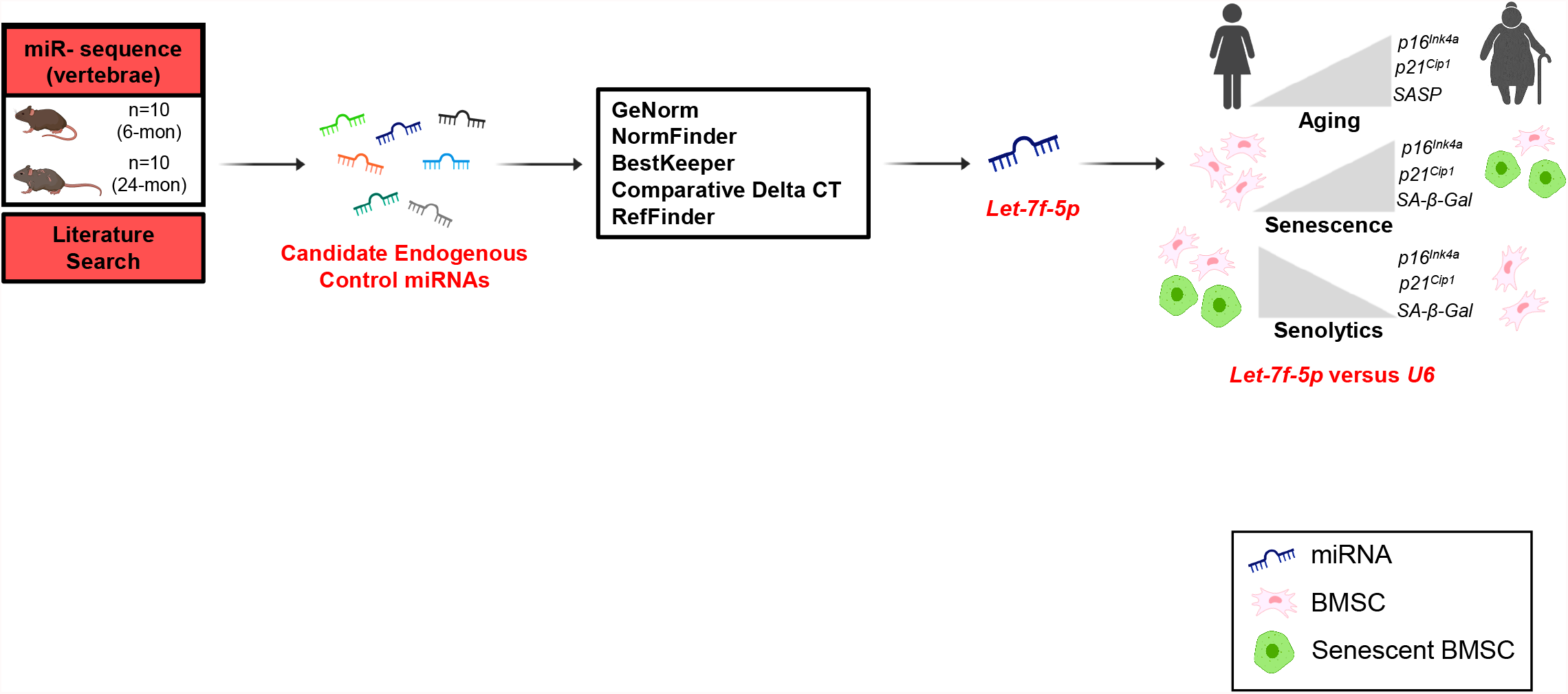
Schematic diagram of the study design. (designed using Biorender.com).

We established our average NormFinder stability scores for each iteration and the minimum score representing the best set of miRNAs for a particular size (Fig. 2a). For our dataset, even a single miRNA had a low minimal stability score (<0.1) and the addition of a second miRNA barely lowered this score further, although the addition of a third miRNA lowered the score to 0.00. Thus, based on these results we speculated that we need anywhere between 1-3 control miRNAs to normalize our dataset; however, we conservatively selected 3 miRNAs as candidate endogenous controls for initial validation. Due to the relatively low read counts of many of these miRNAs, we focused our selection on miRNAs with high normalized counts and relatively low stability scores. Using these criteria, we selected *Let-7i-5p* (avg. normalized counts=87679.671; stability score=0.07), *Let-7f*-5p (avg. normalized counts=47232.569; stability score=0.16) and *miR-185-5p* (avg. normalized counts=971.411; stability score=0.09) as our candidate endogenous control miRNAs.

**Figure 2.**
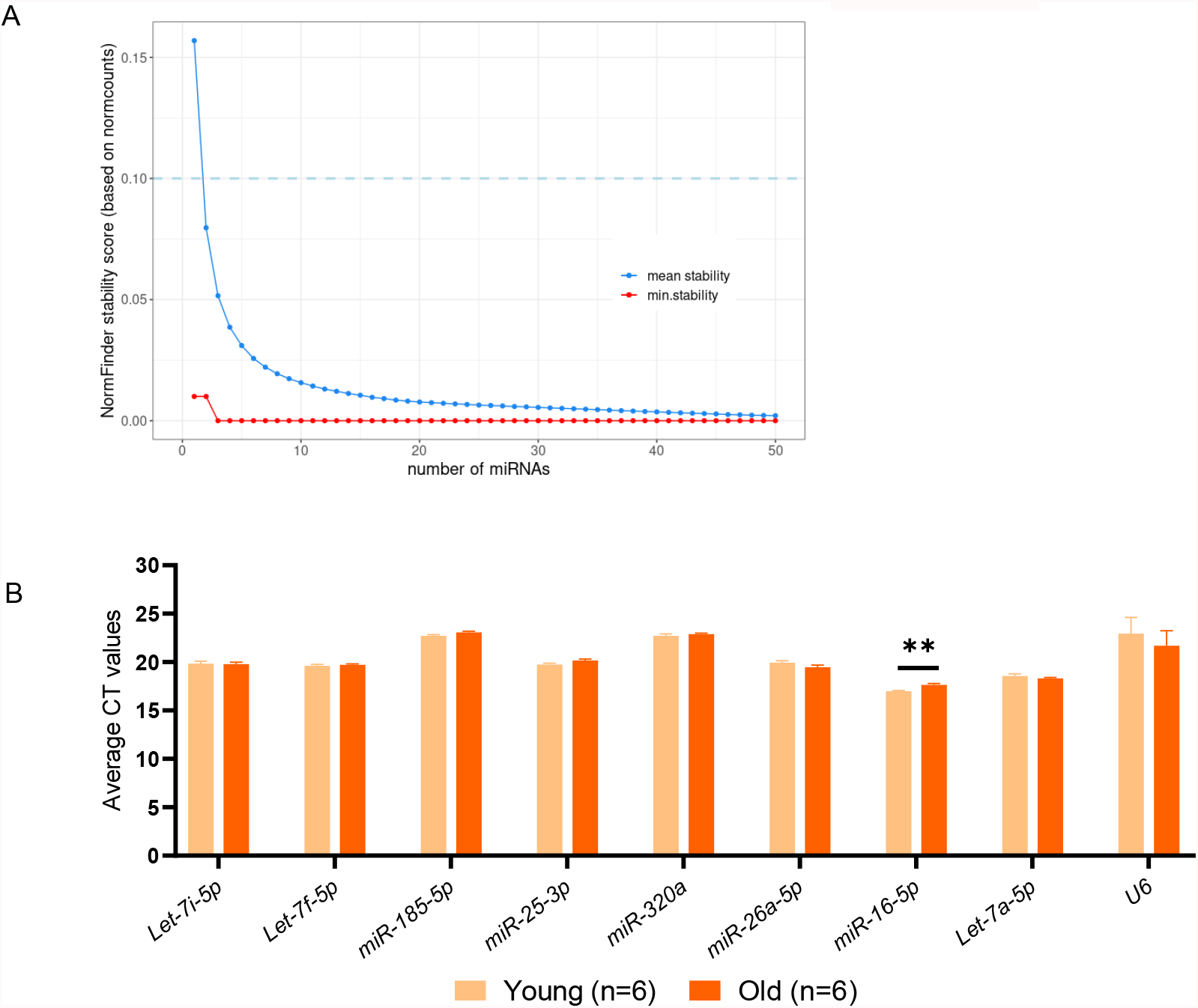
Selection of candidate endogenous control miRNAs. (A) NormFinder mean and minimal stability scores relative to the number of miRNAs iterated. The plot has been limited to first 50 iterations for visualization, beyond which no definite changes were observed. (B) RT–qPCR analysis of the candidate endogenous control miRs and *U6* in the vertebrae of young (6-month) and old (24-month) male and female mice (n=6/group). Data represent mean ± SEM. *p < 0.05; **p < 0.01; ***p < 0.001 (independent samples t-test).

To widen the scope of our study, we performed an extensive literature search for additional miRNAs that were reported to be stable in different biological systems and conditions and incorporated those into our analyses. Our final set of candidate endogenous control miRNAs consisted of: *Let-7i, Let-7f* and *miR- 185-5p* selected from miRNA sequence dataset and *miR-25-3p* (Chen et al., 2016; Drobna et al., 2018; Niu et al., 2016), *-320a* (Allen-Rhoades et al., 2015; Schlosser et al., 2015), *-26a-5p* (Y. Li et al., 2015), *- 16-5p* (Drobna et al., 2018; Song et al., 2012; Wang et al., 2015), and *Let-7a-5p* (Drobna et al., 2018) from the published literature, and each of these were compared to *U6* for their stability (Table 1).

**Table 1.** Candidate endogenous control miRNAs.

| **miRNA** | **Normalized Count** | **NormFinder Stability Score** | **References** |
| --- | --- | --- | --- |
| *Let-7i-5p* * | 87679.67 | 0.07 | - |
| *Let-7f-5p* * | 47232.57 | 0.16 | - |
| *miR-185-5p* * | 971.41 | 0.09 | - |
| *miR-25-3p*† | 10084.71 | 0.21 | Chen et al., 2016; Drobna et al., 2018; Niu et al., 2016 |
| *miR-320a*†# | - | - | Allen-Rhoades et al., 2015; Schlosser et al., 2015 |
| *miR-26a-5p*† | 21518.389 | 0.11 | Y. Li et al., 2015 |
| *miR-16-5p*† | 1712.7345 | 0.27 | Drobna et al., 2018; Song et al., 2012; Wang et al., 2015 |
| *Let-7a-5p*† | 8940.0595 | 0.27 | Drobna et al., 2018 |

\*Selected from miRNA sequence data based on normalized counts and stability scores. † Selected from literature search. The normalized counts and stability scores based on miRNA sequence data are mentioned for reference. # miRNA not expressed in miRNA sequence data.

### Evaluating the stability of selected endogenous control miRNAs with qPCR

We next tested the stability of these candidate endogenous control miRNAs in the osteocyte-enriched samples of young (6-month) and old (24-month) male and female mice (n=6 per group (equal number of males and females)). As shown in Figure 2b, the average of the raw CT values of all miRNAs remains unchanged with aging (p>0.05) except for *miR-16-5p* which increased significantly with aging, and therefore was eliminated from further analyses.

### Selection of *Let-7f* as the endogenous control miRNA in aging

#### To select a stable endogenous control, we used four algorithms

GeNorm, NormFinder, BestKeeper, and comparative Delta CT, available publicly at https://www.heartcure.com.au/reffinder/ which ranked our candidate endogenous controls as per their stability. Finally, the RefFinder tool was used to generate a cumulative stability ranking for these miRNAs based on their average stability calculated by each of these four algorithms, with a lower ranking indicating a higher stability.

#### GeNorm

This algorithm ranked the selected miRNAs in order of their decreasing stability (M-value) with 1.5 considered as the upper limit for a stable miRNA. Although in our analyses all the miRNAs had an M-value of less than 1.5, *miR-185-5p* exhibited the highest stability whereas *U6* was the least stable (Fig. 3a).

**Figure 3.**
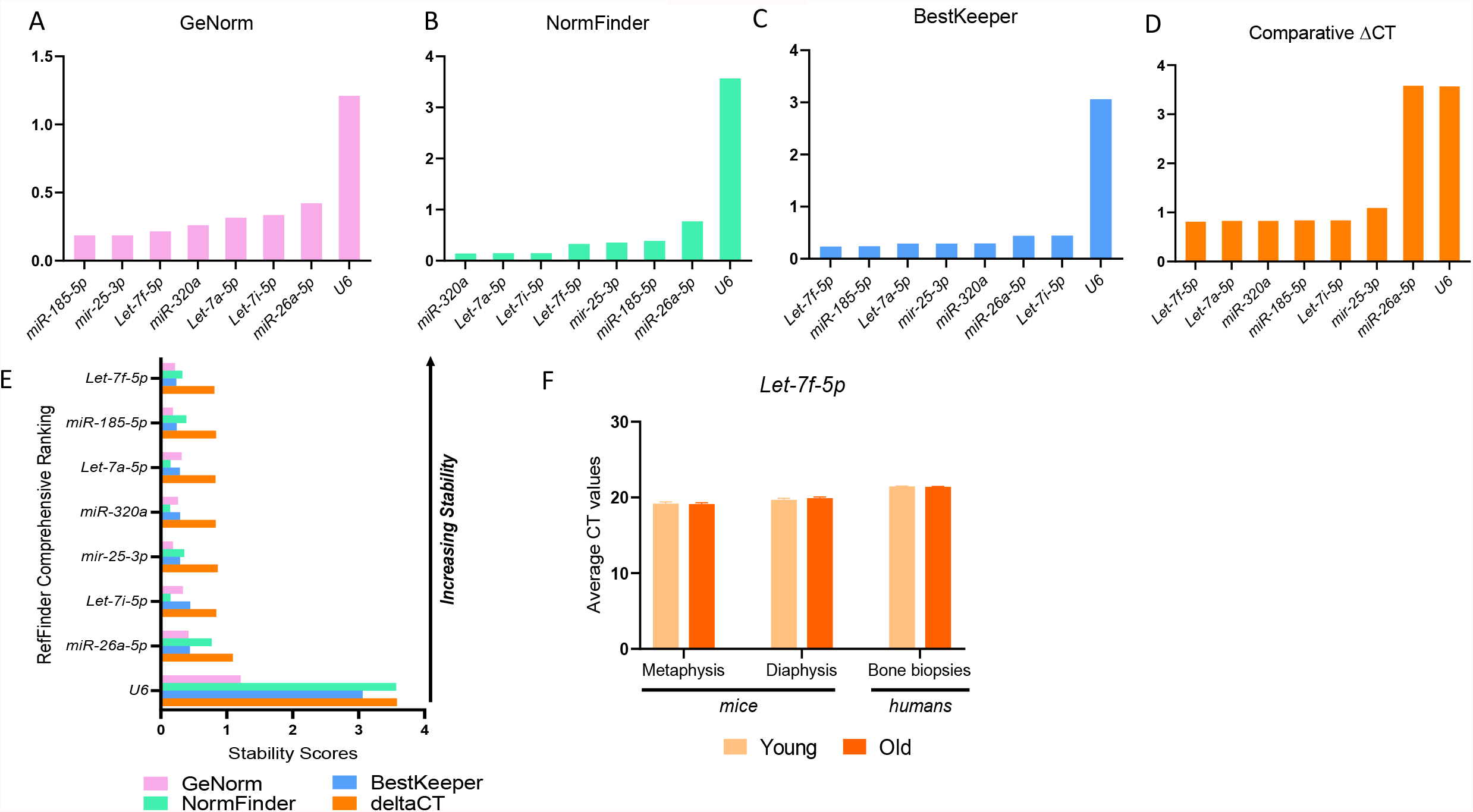
*Let-7f* is the most stable miRNA with aging in bone tissue of mice and humans. Stability ranking of candidate endogenous control miRNAs by (A) GeNorm, (B) NormFinder, (C) BestKeeper, (D) Comparative Delta CT, and (E) RefFinder. (F) RT–qPCR analysis of the *Let-7f* in femur metaphysis and diaphysis of young (6-month) and old (24-month) male and female mice (n=6/group) and in small needle bone biopsies obtained from the posterior iliac crest of 10 young (mean ± SD; 27 ± 3 years) and 10 old (mean ± SD; 78 ± 6 years) healthy female volunteers. Data represent mean ± SEM. *p < 0.05; **p < 0.01; ***p < 0.001 (independent samples t-test).

#### NormFinder

This algorithm uses variation between and within the groups to assess the stability of the candidate miRNAs. It ranked *miR-320a* as the most stable and *U6* as the least stable endogenous control (Fig. 3b).

#### BestKeeper

As per this algorithm, *Let-7f* was the most stable control miRNA. Moreover, the SD value of *U6* (SD=3.06) surpassed the SD threshold (SD=1.0) utilized by this algorithm and was ranked as least stable (Fig. 3c).

#### Comparative Delta CT

Consistent with the results of BestKeeper, this algorithm ranked *Let-7f* as the most stable and *U6* as the least stable gene (Fig. 3d).

#### RefFinder

This algorithm created a final comprehensive stability ranking for the selected miRNAs based on the divergent rankings produced by each of the four algorithms using their geometric mean values.

Based on this analysis, *Let-7f* was the most stable and *U6* remained the least stable endogenous control (Fig. 3e).

Based on these results and those of iterative analyses indicating the sufficiency of using a single miRNA as an endogenous control for our dataset, we selected *Let-7f-5p* as our endogenous control miRNA and validated its stability with aging and senescence in bone tissue and cells.

### Validation of *Let-7f* as a suitable endogenous control in aging bone tissue of mice and humans

We next tested the stability of *Let-7f* at the metaphyses and diaphyses of the femur in young (6-month) and aged (24-month) male and female mice (n=3/group/sex). Consistent with the results at the vertebrae (trabecular bone site), *Let-7f* was stable with aging across both predominantly cortical bone sites.

Moreover, *Let-7f* was also stable in needle bone biopsies (containing both trabecular and cortical bone as well as bone marrow elements) from the posterior iliac crest of young (mean ± SD=27 ± 3 years) and old (mean ± SD=78 ± 6 years) healthy women (n=10/group) who had significantly higher expression of senescence-related genes with aging (Fig. 3f).

### Validation of *Let-7f* as a suitable endogenous control in BMSCs following induction of senescence and senolytic treatment

The stability of *Let-7f* was also tested in non-senescent (vehicle-treated) and senescent (etoposide-treated) BMSCs as well as in senescent BMSCs treated with the senolytic cocktail - dasatinib (D; an FDA-approved tyrosine kinase inhibitor) plus quercetin (Q; a flavanol present in many fruits and vegetables). Senescence was induced in BMSCs using etoposide, a chemotherapeutic agent capable of causing DNA damage, and was confirmed with an increased mRNA expression of *p16*^*ink4a*^ (28.5-fold; p<0.001) and *p21* (8.3-fold; p<0.001) and increased SA-β-Gal activity compared to the vehicle-treated cells (Fig. 4a, b). Moreover, treatment of senescent BMSCs with D+Q was followed by downregulation of *p16*^*ink4a*^ (0.2-fold; p<0.001) and *p21* (0.5-fold; p<0.001) gene expression and fewer SA-β-Gal positive cells compared to the vehicle-treated cells, thus confirming the senolytic effects of D+Q (Fig. 4d, e). *Let-7f* remained stable both with senescence and D+Q treatment in BMSCs (Fig. 4c, f).

**Figure 4.**
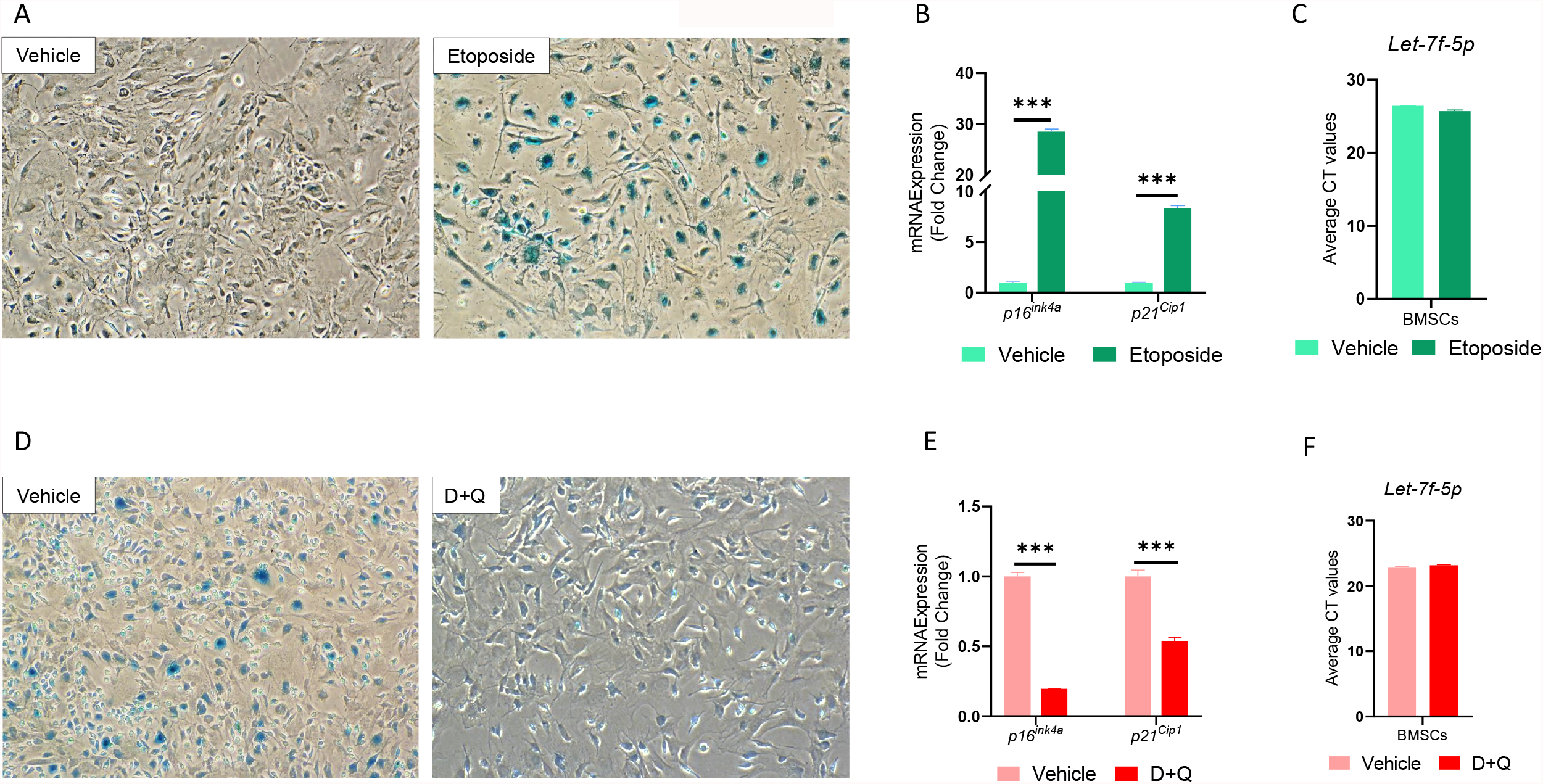
*Let-7f* remains stable in BMSCs following induction of senescence and with senolytic treatment. (A) Representative images of the SA-β-Gal stained BMSCs treated with vehicle (DMSO) and Etoposide (20uM) to induce senescence (magnification, 10X; n = 3/group). RT–qPCR analysis of (B) *p16*^*ink4a*^ and *p21*^*Cip1*^, and (C) *Let-7f-5p*, in BMSCs treated with vehicle (DMSO) and Etoposide (20uM); gene expression was denoted as fold-change relative to vehicle (n = 3/group). (D) Representative images of the SA-β-Gal stained BMSCs treated with vehicle (DMSO) and D and Q to eliminate senescent cells (magnification, 10X; n = 3/group). RT–qPCR analysis of (E) *p16*^*ink4a*^ and *p21*^*Cip1*^, and (F) *Let-7f-5p*, in BMSCs treated with vehicle (DMSO) and D and Q; gene expression was denoted as fold-change relative to vehicle (n = 3/group). Data represent mean ± SEM. *p < 0.05; **p < 0.01; ***p < 0.001 (independent samples t-test).

### Testing the translational application of *Let-7f* versus *U6* for data normalization in humans

To test the application scope of *Let-7f* as a control miRNA versus *U6*, we measured the expression of *miR-34a-5p* in human bone biopsy samples from young vs old women and in BMSCs following induction of senescence and after administration of senolytics (D+Q). When *miR-34a* was measured using *Let-7f* as the endogenous normalizer, its expression increased with aging in human bone biopsies (Fig. 5a; 1.6- fold; p=0.001) and with senescence in BMSCs (Fig. 5b; 1.5-fold; p=0.032) and decreased following treatment with D+Q (Fig. 5c; 0.5-fold; p=0.03). However, when *U6* was used as the endogenous control, *miR-34a* did not change significantly with aging (Fig. 5a; 0.1-fold; p=0.323) and senescence (Fig. 5b; 0.9- fold; p=0.91) and showed an upregulated trend following treatment with D+Q (Fig. 5c; 6.2-fold; p=0.08).

**Figure 5.**
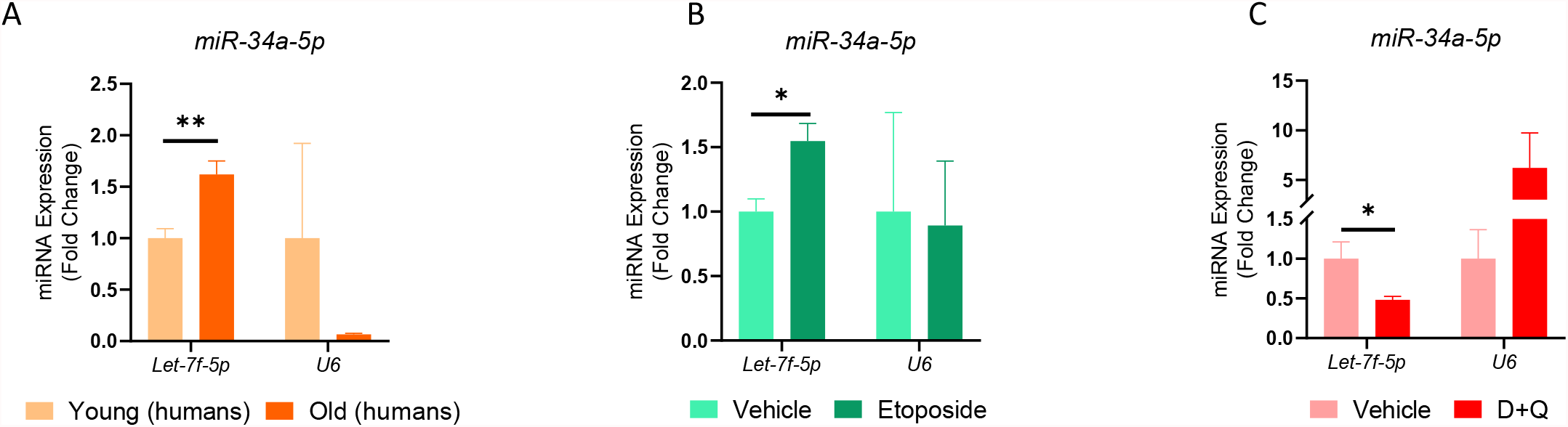
Applicability of *Let-7f* as a control miRNA compared to *U6*. RT-qPCR analyses performed for *miR-34a-5p* using *Let-7f* and *U6* as endogenous control in (A) small needle bone biopsies obtained from the posterior iliac crest of 10 young (mean ± SD; 27 ± 3 years) and 10 old (mean ± SD; 78 ± 6 years) healthy female volunteers, (B) BMSCs treated with vehicle (DMSO) and Etoposide (20uM), and (C) BMSCs treated with vehicle and D and Q; gene expression was denoted as fold-change relative to vehicle (n = 3/group). Data represent mean ± SEM. *p < 0.05; **p < 0.01; ***p < 0.001 (independent samples t-test).

## Discussion

Aging is characterized by a gradual physiological decline resulting in impairment of physical function and increased risk of mortality. Cellular senescence is one of the *‘hallmarks of aging’* that significantly contributes to the aging process and its related phenotype (López-Otín et al., 2013). Multiple studies have shown that senescent cells accumulate with aging in various tissues, including bone and various marrow cell populations, subsequently leading to osteoporosis and its related fractures (Farr et al., 2016, 2017). MicroRNAs have been identified as critical regulators of bone remodeling, aging, and cellular senescence, that mostly downregulate gene expression by binding to the 3’-UTR of the target mRNAs (Bushati & Cohen, 2007; Pignolo et al., 2021). Considering the rising interest in miRNAs as biomarkers, diagnostic and therapeutic agents, and their mechanistic role in the regulation of various biological processes and diseases, inter-experimental reproducibility and comparability of their qPCR-based expression data across various studies is crucial. Thus, the selection of an appropriate endogenous control becomes important for data normalization to generate reliable, comparable, and reproducible results.

In this study, we selected a set of candidate endogenous control miRNAs from our aging-related miRNA sequence dataset and literature search and compared them to *U6*, a widely used control gene in miRNA- related studies. These selected candidate control miRNAs were then validated and ranked for their stability based on four algorithms, GeNorm, NormFinder, BestKeeper, and Comparative Delta CT, and finally, the RefFinder tool created a final stability ranking using their geometric mean values. Although each algorithm established a different miRNA as ‘most stable’ (GeNorm: *miR-185*; NormFinder: *miR- 320a*; BestKeeper and Delta CT: *Let-7f*), *U6* remained the least stable control gene across all algorithms. Our results depicting the instability of *U6* are consistent with previous studies where *U6* is regarded as a highly unstable gene in various conditions (Lou et al., 2015; Xiang et al., 2014). Based on our analyses, we selected *Let-7f* as our endogenous control miR for further validation.

*Let-7f* remained stable at the metaphysis and diaphysis of the femur in mice and in human bone biopsy samples from young versus old women with significantly higher mRNA expression of *p16*^*ink4a*^ and *p21*, as well as a subset of SASP markers. These results indicated the extended applicability of *Let-7f* as a control miRNA in senescence-related conditions. We further reported that *Let-7f* remained stable in BMSCs following induction of senescence with etoposide and in senescent BMSC after treatment with a senolytic cocktail (D+Q), thus expanding the scope of utility of *Let-7f* as a control miRNA to both bone aging, senescence, and monitoring of the selective elimination of senescent cells – *i*.*e*., “senolysis”. We then tested the utility of *Let-7f* as a normalizer in comparison to *U6* by using each to normalize the expression of *miR-34a*, a miRNA well established in the literature to increase with aging and senescence in various cells and tissues. The expression of *miR-34a* was measured in young versus old human bone biopsy samples, and in BMSCs following induction of senescence and treatment with senolytics.

Interestingly, normalization using *Let-7f* resulted in an expected and significant increase in *miR-34a* expression with aging and senescence and a decrease in its expression after treatment with D+Q. However, following normalization to *U6*, the expression pattern of *miR-34a* changed where it decreased with aging and senescence and increased in response to senolytic treatment. Although the fold-changes in each of these conditions were quite large, particularly with aging and senolytic treatment, the results showed no statistical significance owing to the large intra- and inter-group variability. These results are reflective of the highly unstable nature of *U6* and the corresponding misinterpretation of data that can occur on using an unsuitable endogenous normalizer.

Most studies evaluating miRNAs in bone use *U6* or another snRNA as a control, whereas few studies have investigated condition-specific endogenous control miRNAs in bone tissue or cells. Chen et al. reported *miR-25-3p* as the most stable miRNA in serum samples from osteoporotic rats, monkeys and women, and with osteoblast and osteoclast differentiation (Chen et al., 2016). Additionally, *miRs-103a-3p* and *-22-5p* have been reported to be highly stable in extracellular vesicles released from cartilage cells, subcutaneous adipose tissue and BMSCs of osteoarthritic patients and *miR-191-5p* is shown to have high stability in human bone marrow stromal cell lines and primary cells (Costé & Rouleux-Bonnin, 2020; Ragni et al., 2021). The results of each of these studies are unique and different from our results due to differences in methodologies, and the disease states and conditions being tested. Although the selection of our candidate control miRNAs was partly based on miRNA sequence data, we validated our results using qPCR, which is essential as the operating principles used for quantification by the two methods are different. MicroRNA sequencing is based on raw read counts of each transcript whereas qPCR is based on hybridization and amplification of primers and probes. Moreover, qPCR remains the most widely accepted and used method for quantifying gene expression to date (Livak & Schmittgen, 2001; Rajeevan et al., 2001). Furthermore, our results using *Let-7f* or *U6* to measure the change in expression of *miR- 34a* under various conditions re-emphasized our notion that an unstable endogenous control can lead to faulty findings and misinterpretation of results thereby hindering the reproducibility of the study and its comparison across the literature.

A limitation of this study is that although we validated *Let-7f* in BMSCs following induction of senescence and senolytic treatment, our miRNA sequence data was derived from young and aged mice. It has been shown previously that senescent cells accumulate with aging; however, we acknowledge that our aged bone tissue samples were not a pure senescent cell population. For our study design, the ideal selection of candidate control miRNAs should be made from three different miRNA sequence datasets – young versus aged, non-senescent vs senescent BMSCs, and vehicle versus senolytic treated BMSCs.

However, conducting the experiments at such a vast scale was beyond the scope of the current study, which opens the possibility that another miRNA by itself or in combination with *Let-7f* may be as or even more stable than *Let-7f* alone. Another limitation is that we were not able to compare our endogenous control miRNA other small RNAs such as SNORD44, SNORD48, SNORD68, SNORD116 etc., that have been used as control genes for miRNA related studies (Bignotti et al., 2016; Masè et al., 2017). Although like U6, these small RNAs are structurally different from miRNAs which is a caveat for their use as an endogenous control. Therefore, future studies will be needed to assess the stability of these and other small RNAs.

In conclusion, we used a systematic approach to identify *Let-7f* as a suitable endogenous control miRNA that remains stable in the bone tissue of mice and humans with aging and in BMSCs with senescence and in response to senolytic treatment and confirmed its translational applicability by using it as a normalizer to assess the expression of a well-established aging and senescence-associated miRNA.

## HIGHLIGHTS

- *Let-7f* is the most stable miRNA with aging at cortical and trabecular bone sites in young and old mice, and in human bone tissue samples from young and old women with high levels of senescence and SASP related genes.
- *Let-7f* maintains its stability in BMSCs following induction of senescence and in senescent BMSCs after treatment with senolytic cocktail of dasatinib plus quercetin.
- When *Let-7f* is used as a normalizer versus *U6, miR-34a* expression increases with aging in human bone tissue samples and in BMSCs from mice following induction of senescence and decreases in senescent BMSCs after treatment with dasatinib plus quercetin.

## Notes

### Competing Interest Statement

The authors have declared no competing interest.

